# Beyond Antimicrobial Activity: Soil Bacteria Reveal a Biotransformation Fate for the Lanthipeptide Nisin

**DOI:** 10.64898/2026.07.06.736734

**Authors:** Quynh Khoa Pham, Carlos N. Lozano-Andrade, Kah Yean Lum, Mikael Lenz Strube, Lars Jelsbak, Thomas O. Larsen, Scott A. Jarmusch

**Affiliations:** Department of Biotechnology and Biomedicine, Technical University of Denmark, Søltofts Plads 221, Kgs. Lyngby DK-2800, Denmark

**Author notes:** Address correspondence to Scott A. Jarmusch.

**Keywords:** nisin, biotransformation, *Pseudomonas fragi*, *Burkholderia stabilis*, soil microbiome, nutrient recycling, LC-MS

## Abstract

Natural products are central mediators of microbial interactions. However, once released into the environment, they also become available for neighboring microorganisms capable of degrading and modifying them through biotransformation. These biotransformations may fundamentally reshape metabolomes and influence community behavior, yet our understanding of these processes remains limited. Ribosomally synthesized peptides are particularly compelling in this context because their structural complexity and potent antimicrobial activity coexist with the potential to yield essential nutrients and reduced bioactivity through biotransformation. Identifying the pathways underlying these biotransformations is essential for understanding mechanisms that support microbial coexistence and nutrient recycling in soil microbiomes. Here, we used nisin as a model peptide to investigate biotransformation by soil bacteria. Selective isolation under nisin-rich, carbon-limited conditions yielded two Gram-negative isolates, *Burkholderia stabilis* and *Pseudomonas fragi*. Using growth assays and liquid chromatography–mass spectrometry, we found that both isolates grow in the presence of nisin while biotransforming and depleting the peptide. *Burkholderia stabilis* completely converted nisin through sequential cleavage of the C-terminus, hinge region and lanthionine ring C, whereas *Pseudomonas fragi* showed more limited processing restricted to the C-terminal region. Although these biotransformations dismantled structural features required for nisin’s antimicrobial activity, the intrinsic resistance of both isolates suggests a role beyond detoxification. We further detected nisin biosynthetic genes in the source environment, supporting nisin’s ecological relevance and suggesting that these bacteria may participate in its turnover in soil. Together, these findings reveal extensive microbial processing of nisin and support a role for antimicrobial peptide recycling in soil microbiomes.

**Importance:** Natural products are often studied through the lens of their biological activities, but much less attention has been paid to what happens to these molecules after they enter complex microbial communities. Using the lantibiotic nisin as a model system, we show that soil bacteria can extensively biotransform and deplete an antimicrobial peptide through extracellular enzymatic activity. The presence of both nisin-producing and nisin-biotransforming microorganisms in the same soil environment suggests that antimicrobial peptides may be continuously produced and recycled in nature. Our findings highlight biotransformation as an important but underexplored process governing the persistence, turnover, and ecological roles of microbial natural products.

## Introduction

Natural products are central mediators of microbial interactions, shaping competition, cooperation, and resource acquisition within complex communities (1–5). In soil ecosystems, among the most taxonomically and metabolically diverse habitats on Earth (6–10), these molecules do not persist as static chemical entities but instead become integrated into dynamic metabolic networks. Consequently, natural products can be modified, degraded, or repurposed by neighboring microorganisms through biotransformation processes. Previous studies have shown that such transformations can contribute to detoxification and resistance (11) or enable the use of natural products as nutrient sources (12, 13), with important consequences for microbial interactions and community structure (14). However, these observations derive from a limited number of systems, and the broader ecological significance of natural product biotransformation in soil microbiomes remains unclear. In particular, the microorganisms involved, the pathways they employ, and the contribution of these processes to natural product turnover in soil environments remain poorly understood.

Ribosomally synthesized and post-translationally modified peptides (RiPPs) are a widespread class of natural products in soil ecosystems (15). Their precursor peptides are enzymatically modified to generate structurally diverse molecules that often display potent antimicrobial activity (16–18). At the same time, their extensive post-translational modifications often enhance structural stability and protect them from proteolytic degradation (19). Nevertheless, some microorganisms can enzymatically biotransform these molecules, resulting in a marked loss of biological activity (20). Despite these potentially important ecological consequences, the microorganisms responsible for RiPP biotransformation and the pathways by which these compounds are biotransformed in natural environments remain largely unknown.

Among RiPPs, the lantibiotic nisin is one of the best-characterized members of the class. Produced by certain strains of *Lactococcus lactis*, nisin is a 34-amino-acid peptide containing multiple dehydrated residues and five lanthionine rings that confer structural stability and potent antimicrobial activity against Gram-positive bacteria (21). Nisin Z and nisin A are the most widespread natural variants and differ by a single amino acid, asparagine and histidine, respectively, resulting in similar antimicrobial activity but distinct physicochemical properties (21–23). Of which, nisin Z is more widely distributed among nisin producers than nisin A (24). Because of its well-defined structure, established biosynthetic pathway, broad distribution, and commercial availability, nisin Z provides an attractive model system for investigating RiPP biotransformation. Despite its structural complexity and resistance to proteolytic degradation, nisin biotransformation has been reported in a limited number of food-and host-associated bacteria (20, 25). The best-characterized example is cleavage of the C-terminus by the nisin resistance protein (NSR), which reduces antimicrobial activity and contributes to resistance (20). However, little is known about the microorganisms capable of biotransforming nisin in soil environments, the pathways by which these biotransformations occur, or their ecological consequences.

In this study, we investigated the biotransformation of nisin Z as a model lanthipeptide within soil microbiomes. We identified soil bacteria capable of growing in the presence of nisin and demonstrated extensive biotransformation of the peptide, including the formation of previously undescribed biotransformation products and pathways. We further provide evidence that extracellular proteases mediate these biotransformations and show that nisin-producing populations coexist with nisin-biotransforming bacteria within the same soil environment. Together, these findings reveal an underexplored route of antimicrobial peptide turnover in soil microbiomes and suggest that biotransformation contributes not only to the chemical fate of RiPPs but also to nutrient recycling within microbial communities.

## Results

### Soil-derived Pseudomonas fragi and Burkholderia stabilis subsist on nisin

To enrich for bacteria capable of biotransforming nisin, soil from site P9 (grassland soil from the Jægersborg Deer Park, a UNESCO World Heritage Site) was suspended in PBS and inoculated into M9 minimal medium containing 0.01% glucose and 0.01% nisin. After seven days of incubation, the medium became turbid, indicating microbial growth under these low-nutrient conditions. The diluted culture was plated onto M9 agar supplemented with or without nisin. After incubation, the nisin-containing plate displayed more abundant bacterial growth and two distinct colony morphologies compared with the plate lacking nisin (Fig. 1A). Representative colonies were isolated, purified on LB medium, and cryopreserved for further characterization.

**Figure 1.**
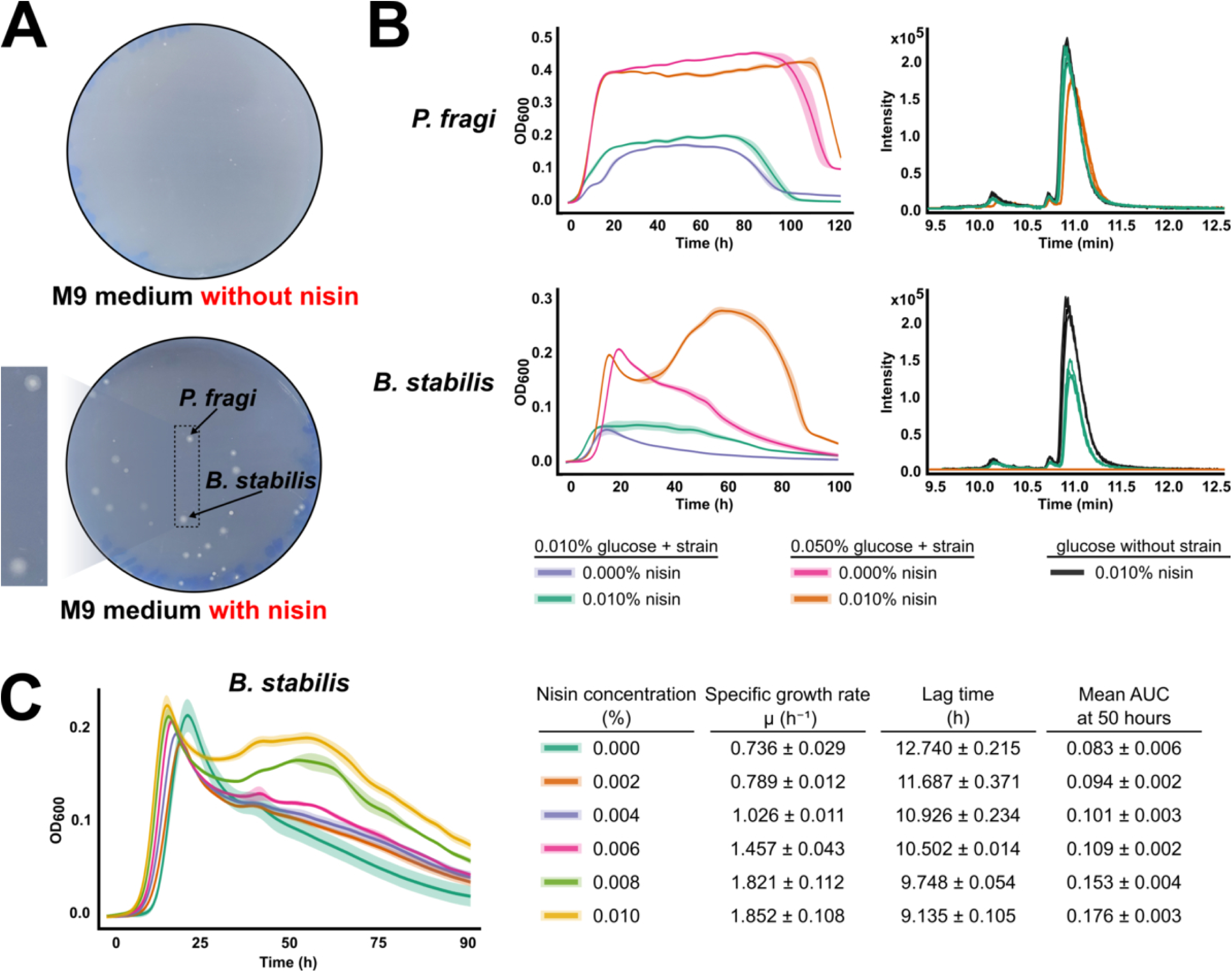
Initial isolation and screening of soil bacteria capable of biotransforming nisin are conducted using M9 minimal medium supplemented with 0.01% nisin. (A) The growth of bacteria is observed on M9 minimal agar plates containing 0.01% glucose, with and without nisin. (B) Growth curves for *P. fragi* and *B. stabilis* in M9 medium broths containing 0.01% and 0.05% glucose, with and without nisin. Extracted ion chromatograms of nisin in these cultures are compared with those in M9 medium broths containing nisin but no bacteria. (C) Nisin concentration-dependent growth curves for *B. stabilis*.

To identify both isolates, their complete genome sequences were obtained using Oxford Nanopore Technologies and analyzed with autoMLST 2.0. The findings revealed that strain 1 exhibited 99.2% average nucleotide identity (ANI) with the genome of *Pseudomonas fragi* (Fig. S1), and strain 2 demonstrated 98.5% ANI to *Burkholderia stabilis* (Fig. S2).

To evaluate whether the isolates could grow in the presence of nisin and potentially biotransform the peptide, growth assays were performed in microtiter plates using M9 minimal medium supplemented with 0.01% glucose and with or without 0.01% nisin (18 µg per well). Growth curves were monitored by optical density (OD_600_) and nisin abundance was measured by LC–MS at the experimental endpoint. *P. fragi* showed improved growth in the presence of nisin, with a shorter lag phase (2.307 ± 0.352 h versus 8.761 ± 0.874 h) and a higher OD_600_ at 12 h (0.112 ± 0.010 versus 0.054 ± 0.001) than cultures grown without nisin. Despite this growth benefit, little depletion of the peptide was observed (Fig. 1B). *B. stabilis* showed enhanced growth in the presence of nisin, with a shorter lag phase (3.229 ± 0.144 h versus 5.586 ± 0.567 h) and a higher OD_600_ at 40 h (0.060 ± 0.008 versus 0.023 ± 0.001) than cultures grown without nisin. Unlike *P. fragi*, this growth benefit was accompanied by approximately 50% depletion of the initial nisin signal (Fig. 1B). Increasing the glucose concentration to 0.05% enhanced nisin utilization by both strains. Under these conditions, *P. fragi* exhibited prolonged growth and depleted approximately 30% of the peptide, whereas *B. stabilis* displayed a biphasic growth pattern and completely depleted nisin by the end of the incubation period (Fig. 1B).

Because *B. stabilis* completely depleted nisin, we further examined its growth response across a range of nisin concentrations (Fig. 1C). Increasing nisin concentrations led to higher growth rates (0.736 ± 0.029 to 1.852 ± 0.108 µ/h) and shorter lag phases (12.740 ± 0.215 to 9.135 ± 0.105 h). The biphasic growth pattern became more pronounced at higher nisin concentrations and less apparent at lower concentrations. Cultures supplemented with higher nisin concentrations also exhibited greater overall growth, as reflected by increased AUC values over the 50 h incubation period.

### *Pseudomonas fragi* biotransforms nisin through C-terminal cleavage

Following growth in M9 minimal medium supplemented with 0.05% glucose, cultures containing *P. fragi* and nisin, nisin-only controls, and *P. fragi*-only controls were collected from the growth assays. Samples were extracted and analyzed by LC-MS to detect nisin biotransformation products, which were observed only in cultures containing both *P. fragi* and nisin. To describe these products, we defined the following nomenclature: N-terminal fragments (N), middle fragments (M), C-terminal fragments (C), and combinations of these for fragments spanning multiple regions (NM).

After incubation of nisin with *P. fragi*, six biotransformation products, NM1 and C6–C2, were detected (Fig. S3). Exact mass analysis (Fig. 2B) showed that NM1 (2695.1694 Da) exhibited a mass loss of 633.3597 Da relative to nisin (3328.5291 Da), corresponding to the removal of the six C-terminal amino acid residues (Fig. 2A). Conversely, C6 (651.3697 Da) (Fig. 2B) matched the expected mass of the released C-terminal hexapeptide (Fig. 2A). The identities of NM1 and C6 are consistent with previously reported nisin degradation products observed in *Lactococcus* spp. (20). In addition, four related products, C5–C2, displayed sequential mass losses of 87.0313, 113.0846, 137.0584, and 99.0684 Da relative to C6 (Fig. 2B), corresponding to the successive removal of S29, I30, H31, and V32 (Fig. 2A). These observations suggested stepwise degradation of the C-terminal fragment following the initial cleavage event. The proposed structures of NM1 and C6–C2 were shown in Fig. 2, with mass errors ranging from −0.88 to 0.60 ppm.

**Figure 2.**
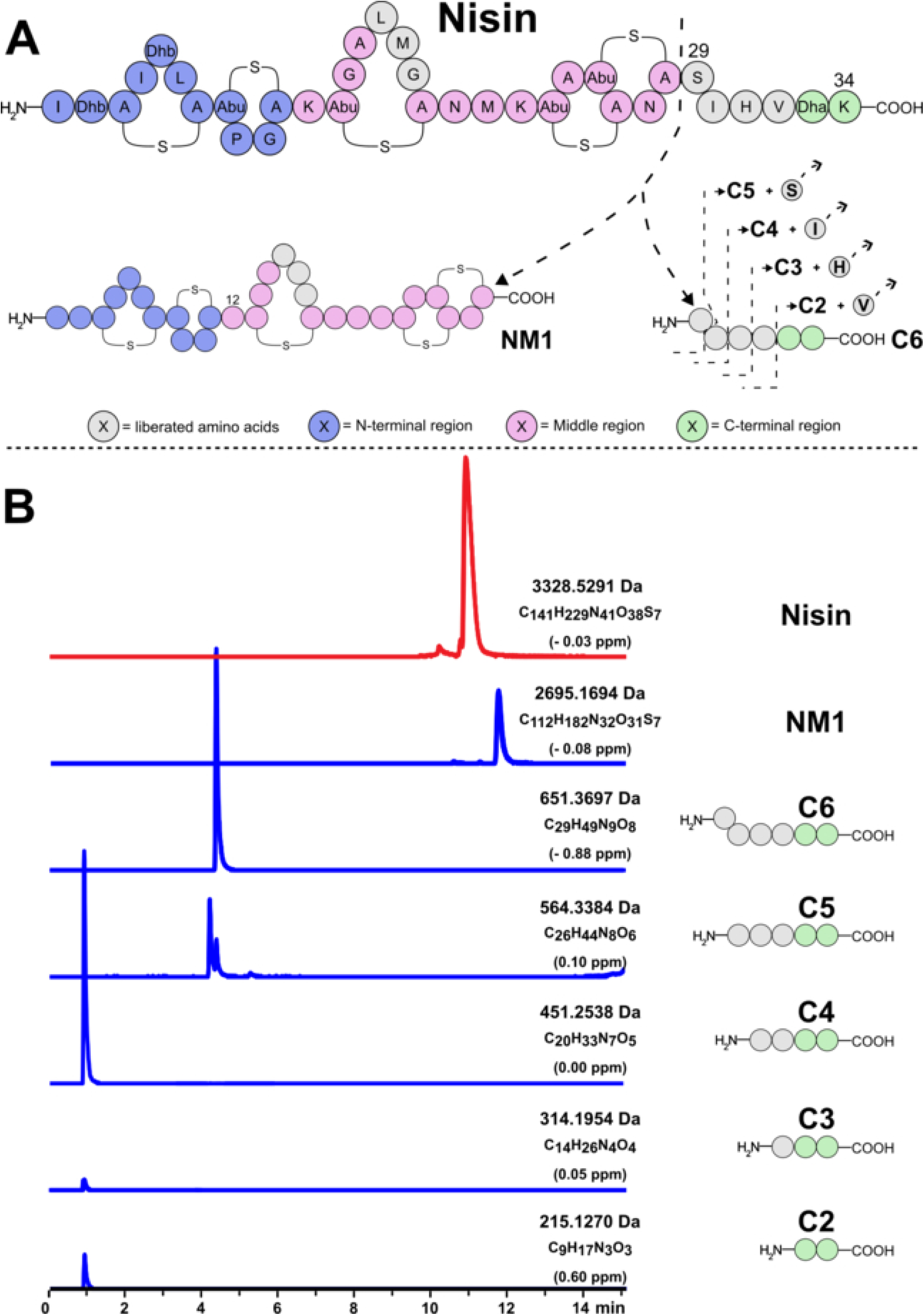
Proposed nisin biotransformation pathways by *P. fragi* (A) and comparison of the retention times and exact masses of nisin and the detected biotransformation products, NM1 and C6–C2 (B). Biotransformation products are detected following incubation of nisin with *P. fragi* in M9 minimal medium supplemented with 0.05% glucose.

### *Burkholderia stabilis* mediates extensive biotransformation of nisin

Similarly to *P. fragi*, nisin biotransformation products generated by *B. stabilis* were identified by LC-MS based on peaks uniquely detected in cultures containing both *B. stabilis* and nisin. After incubation in M9 minimal medium supplemented with 0.05% glucose for 100 h, 15 biotransformation products (NM2–NM6, C5–C2, M1–M5, and N1) were detected (Fig. S4–S6).

Exact mass analysis indicated that NM2 resulted from the removal of the five C-terminal amino acid residues of nisin, while C5 corresponded to the released C-terminal pentapeptide (Fig. 3). In pathway 1, these products were consistent with initial cleavage at the S29–I30 peptide bond, one residue further toward the C-terminus than the cleavage site observed for *P. fragi*. In pathway 2, subsequent stepwise degradation of the released C-terminal fragment generated products C4–C2 through sequential removal of I30, H31, and V32.

**Figure 3.**
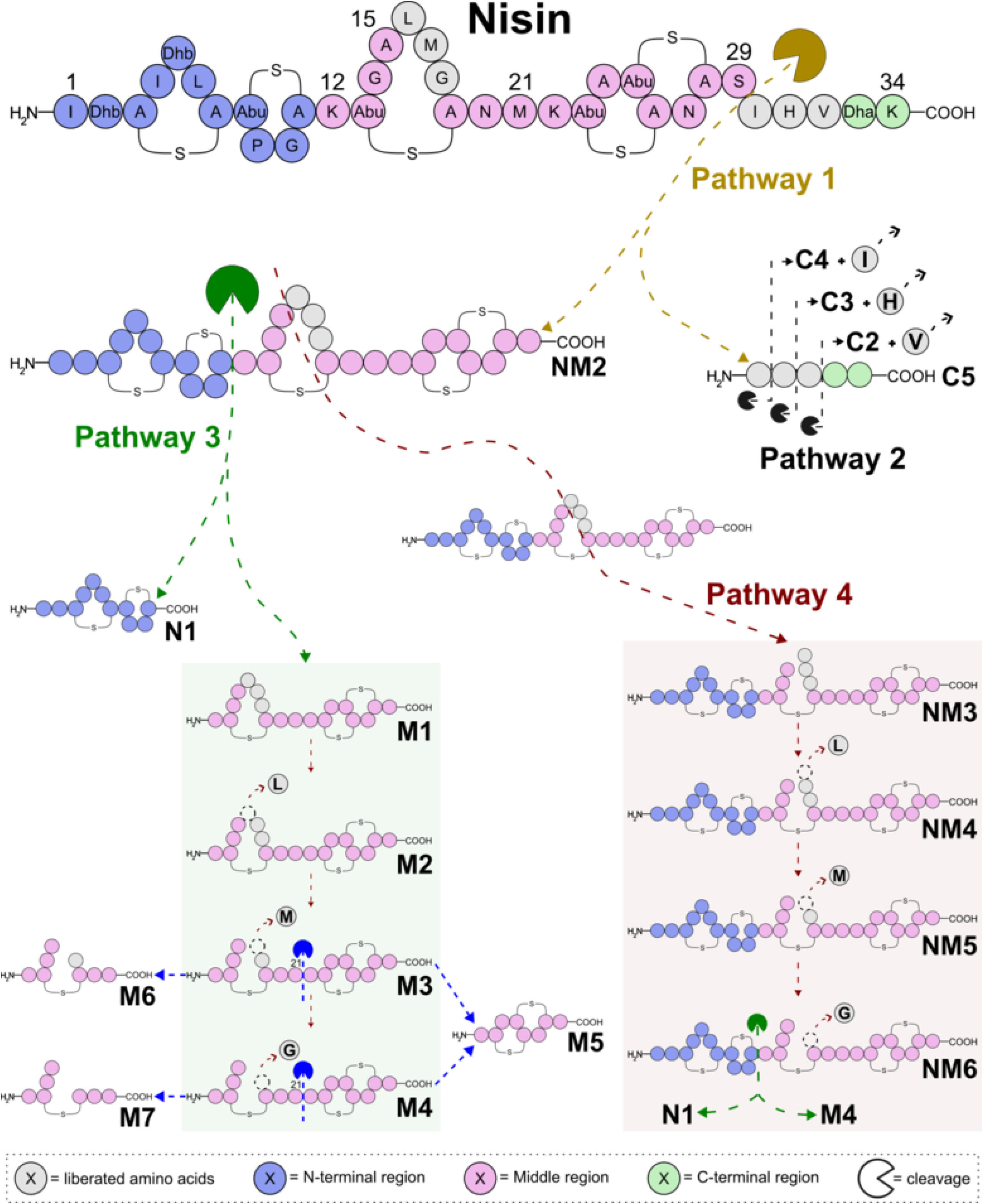
Proposed nisin biotransformation pathways by *Burkholderia stabilis* (pathways 1–4). Biotransformation products are detected following incubation of nisin with *B. stabilis* in M9 minimal medium supplemented with 0.05% glucose for 100 h, except for M6 and M7 which are detected following incubation of nisin in sterile culture supernatants derived from *B. stabilis* grown in KB medium for 70 h.

In addition to C-terminal processing, NM2 underwent further degradation through two interconnected pathways (Fig. 3). In pathway 3, cleavage at the A11–K12 peptide bond generated the stable N-terminal product N1 and middle fragment M1. Successive removal of residues from lanthionine ring C produced intermediates M2–M4, which were subsequently processed to the stable endpoint product M5. In pathway 4, hydrolytic opening of lanthionine ring C generated NM3, followed by sequential removal of residues to produce NM4–NM6. Cleavage of NM6 at the A11–K12 peptide bond generated N1 and M4, linking pathways 3 and 4.

The proposed structures of all biotransformation products were shown in Fig. 3 and were supported by exact mass measurements with mass errors ranging from −1.18 to 1.10 ppm (Fig. S7–S9). The structures of the endpoint products N1, M5, and C2–C3 were further supported by MS/MS fragmentation patterns consistent with the proposed structures and charge-migration fragmentation mechanisms (Fig. S11–S14) (26, 27).

### Extracellular proteases from *Burkholderia stabilis* mediate nisin biotransformation

To assess whether extracellular enzymes from *Burkholderia stabilis* mediated nisin biotransformation, we incubated nisin in cell-free supernatants obtained from cultures grown in either M9 medium supplemented with 0.5% glucose or KB medium. When the supernatants were heat-treated prior to incubation, nisin biotransformation was not observed with LC–MS (Fig. S17). In contrast, nisin was biotransformed when incubated with untreated supernatants (Fig. S17), with the extent of biotransformation varying based on the growth medium (Fig. 4).

**Figure 4.**
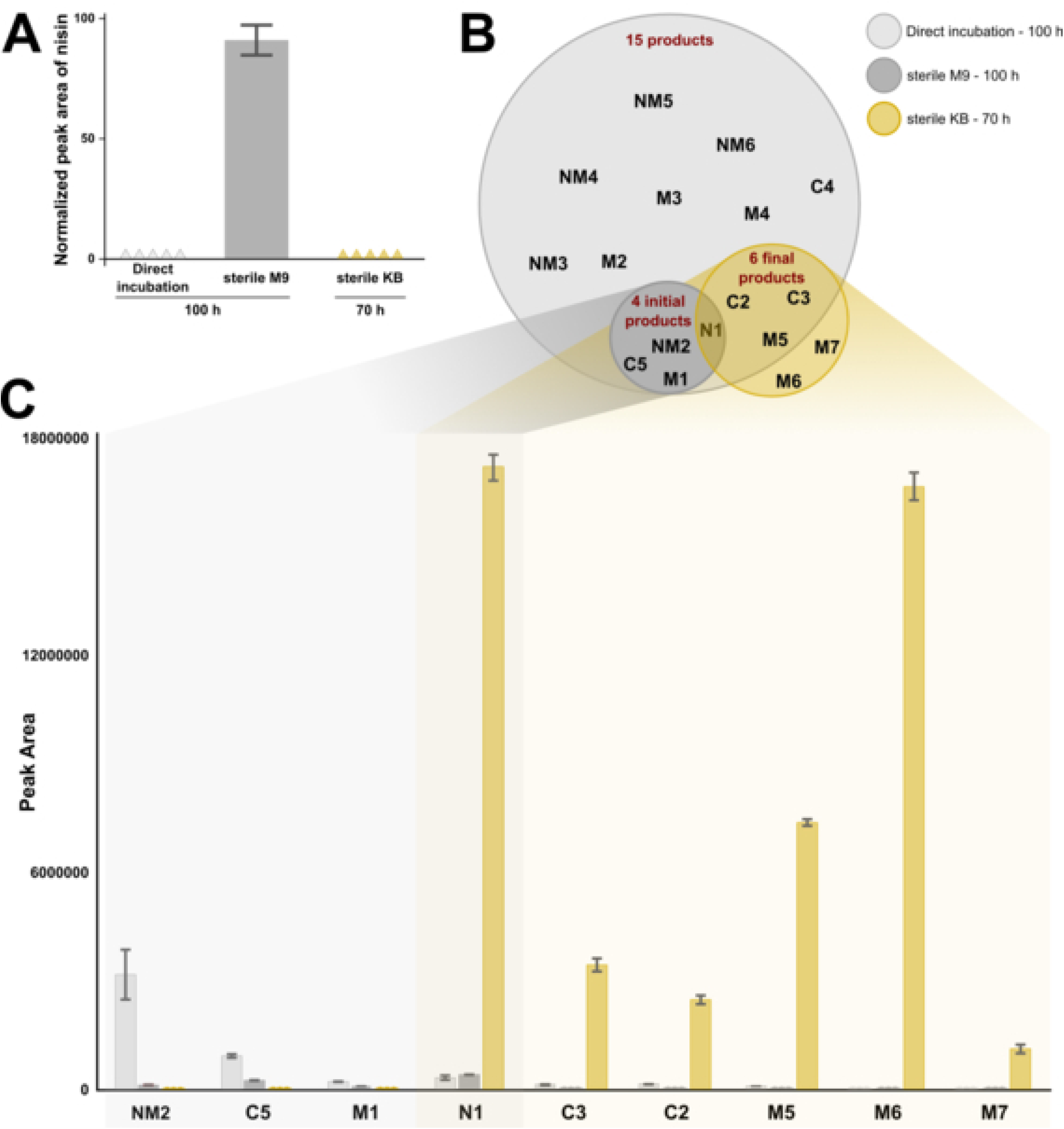
Nisin depletion and biotransformation under different incubation conditions. (A) Nisin depletion following direct incubation with *B. stabilis* cells in M9 medium supplemented with 0.05% glucose for 100 h (Direct incubation), incubation in sterile-filtered supernatants obtained from *B. stabilis* cultures grown in M9 medium supplemented with 0.5% glucose for 100 h (Sterile M9), or incubation in sterile-filtered supernatants obtained from *B. stabilis* cultures grown in KB medium for 70 h (Sterile KB). (B) Detection of nisin biotransformation products under each condition. (C) Comparison of the relative abundances of early-stage and late-stage biotransformation products.

In supernatants from M9 cultures, nisin showed only minor depletion after 100 h (Fig. 4A). Correspondingly, only early intermediates of pathways 1 and 3 (Fig. 3) were detected at low abundance, including NM2, C5, N1, and M1 (Fig. 4B-C). This limited biotransformation occurred despite cultures reaching high cell densities in M9 medium (OD_600_ = 1.79 ± 0.05), suggesting that enzyme production or activity, rather than bacterial growth, constrained nisin biotransformation under these conditions.

Supernatants derived from KB cultures displayed substantially greater nisin biotransformation activity than those derived from M9 cultures. Under these conditions, nisin was completely depleted after 70 h (Fig. 4A), consistent with the biotransformation observed when nisin was incubated directly with *B. stabilis* cells. Notably, cultures grown in KB and M9 media reached similar cell densities (OD600 = 1.87 ± 0.01 and 1.79 ± 0.05, respectively), indicating that the differences in biotransformation activity were not attributable to differences in bacterial growth.

LC-MS analysis revealed that later-stage pathway products, including N1, C2, C3, and M5, accumulated at high abundance in KB-derived supernatants (Fig. 4B-C). In addition, M6 and M7 were detected in KB-derived supernatants but were not observed when nisin was incubated directly with *B. stabilis* in M9 medium. The detection of M6 and M7 provided further support for the proposed pathway 3 biotransformation route (Fig. 3). Proposed structures were supported by retention times, isotope distributions, exact mass measurements, and MS/MS fragmentation patterns (Fig. S10 and Fig. S15-S16).

### Nisin-producing and nisin-biotransforming bacteria coexist within the same soil microbiome

To assess the ecological relevance of nisin biotransformation, we screened for nisin precursor genes, *nisA* and *nisZ*, in soil samples collected from three sites (P5, P9, and P16) within the Dyrehaven. Notably, site P9 was the location where the nisin biotransformers, *P. fragi and B. stabilis*, were isolated. The precursor genes *nisA* and *nisZ* are the two most common nisin variants and differ by a nucleotide at position 148 (C in *nisA* and A in *nisZ*) within the precursor peptide coding region (24) (Fig. 5A). Because structural diversity among RiPPs is primarily encoded within the precursor peptide sequence, primers were designed to amplify a region spanning the *nisA/nisZ* gene and part of *nisB*, while preserving sequence variation within the precursor peptide region.

**Figure 5.**
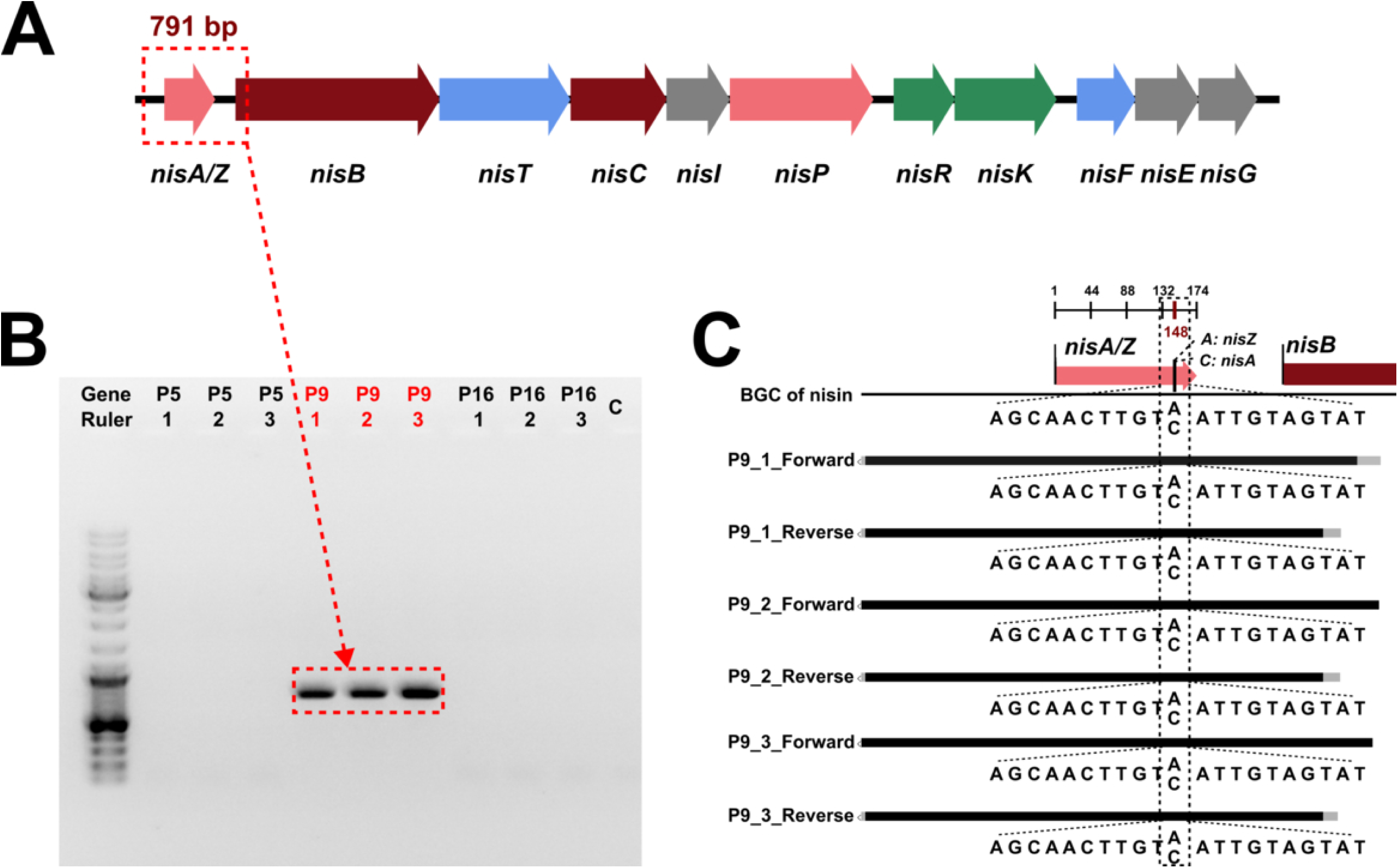
Detection of nisin biosynthetic genes in soil microbiomes. (A) Organization of the nisin biosynthetic gene cluster showing the location of primers used to amplify a 791 bp fragment spanning the *nisA/nisZ* and *nisB* regions. (B) Agarose gel electrophoresis of PCR products generated from environmental DNA extracted from soil samples collected at sites P5, P9, and P16. A ∼791 bp amplicon is detected only in samples from site P9. (C) Sequence analysis of the amplified region showing the presence of both *nisA* and *nisZ* alleles in P9 samples. The nucleotide polymorphism distinguishing *nisA* and *nisZ* is indicated.

Using these primers, we detected a single 791 bp amplicon in all triplicate samples collected from site P9, whereas no products were detected from site P5 or P16 (Fig. 5B). Sequence analysis of the amplified fragments revealed both C and A nucleotide at the diagnostic position distinguishing the two precursor peptides (Fig. 5C). Because most producers contain either *nisA* or *nisZ* only in their gene (28), the detection of both alleles indicated the presence of multiple genetically distinct nisin-producing populations within the P9 soil microbiome. The presence of both nisin biotransformers and nisin producers at a single site highlighted the crucial connection between nisin production and biotransformation in the same environmental context.

## Discussion

In this study, we demonstrate that soil-derived bacteria can survive and proliferate in nisin-rich, carbon-limited environments through a combination of intrinsic resistance and nisin biotransformation. Using a selective enrichment strategy, we isolated two Gram-negative bacteria, *Pseudomonas fragi* and *Burkholderia stabilis*, which exhibited sustained growth in the presence of nisin. Although the outer membrane of Gram-negative bacteria is known to confer intrinsic resistance to nisin and other antimicrobial peptides (29, 30), the concurrent depletion of nisin, extensive biotransformation of the peptide, and dose-dependent growth responses (Figs. 1-3) support the conclusion that both isolates derive nutritional benefit from nisin. The extent of biotransformation varied substantially between species, with *B. stabilis* exhibiting greater activity than *P. fragi*. Biotransformation was also enhanced under conditions of increased carbon availability, indicating that nutrient status influences the efficiency with which environmental bacteria process antimicrobial peptides.

Our findings support the broader concept that antimicrobial compounds can function not only as inhibitors but also as nutrient resources within microbial communities. Although the ability of environmental bacteria to metabolize antibiotics under nutrient limitation has been documented for small-molecule compounds such as β-lactams (31) and aminoglycosides (32), analogous processes involving RiPPs remain largely unexplored. By enriching bacteria capable of growing in the presence of nisin as the dominant available peptide substrate, we uncovered a metabolic strategy that may be more widespread than currently appreciated. The diversity of both antimicrobial peptides and environmental microorganisms suggests that many additional peptide biotransformation pathways remain undiscovered. These observations suggest that the ecological lifespan of antimicrobial peptides extends beyond their initial inhibitory activity, continuing through subsequent biotransformation and nutrient recycling.

A key contribution of this study is the demonstration of extensive in vivo biotransformation of nisin, a structurally complex lanthipeptide. Previous studies have primarily characterized nisin degradation using purified enzymes, including nisin-resistance protein, trypsin, chymotrypsin, and thermolysin, revealing individual cleavage events under controlled conditions (20, 33). In contrast, our findings reveal extensive biotransformation pathways operating within living bacterial systems. While *P. fragi* exhibited relatively limited activity, *B. stabilis* generated a diverse series of intermediates through interconnected biotransformation pathways, yielding 17 previously unreported biotransformation products (Figs. 2-3). These pathways involved progressive peptide bond hydrolysis across multiple regions of the molecule, including the C-terminus, the hinge region and lanthionine ring C. Particularly notable was the observation of biotransformation within lanthionine ring C, a structurally constrained region that contributes to the stability and antimicrobial activity of nisin (25, 33). Although cleavage within this region has previously been observed using purified proteases in vitro (33), our results demonstrate that analogous biotransformations can occur during microbial biotransformation in a biological context. The diversity of the observed intermediates and cleavage products suggests that nisin biotransformation involves multiple enzymatic activities acting on distinct regions of the molecule. Because both isolates are intrinsically resistant to nisin (29, 30), these biotransformation pathways are unlikely to function primarily in detoxification. Instead, they point toward a broader role for nisin biotransformation in microbial nutrient acquisition and recycling.

Our results further indicate that extracellular proteases play a central role in nisin biotransformation by *B. stabilis*, consistent with previous reports of extracellular protease production in this species (34). Nisin biotransformation was observed only when cell-free supernatants retained enzymatic activity, whereas heat treatment abolished biotransformation (Fig. S17), demonstrating that active extracellular enzymes are required for the process. The extent of biotransformation also varied substantially across growth conditions. Supernatants derived from cultures grown in KB medium supported more complete nisin biotransformation than those obtained from M9 medium (Fig. 4), despite similar final cell densities. Moreover, direct incubation of nisin with *B. stabilis* cells in M9 medium resulted in more extensive biotransformation than incubation in sterile M9-derived supernatants, suggesting that M9 medium may be a suitable option for further proteomic studies to screen for enzymes produced by *B. stabilis* that are involved in nisin biotransformation. Together, these observations suggest that environmental conditions influence the production, activity, or composition of extracellular enzymes involved in nisin biotransformation. Genome analyses provided further evidence that the responsible enzymes differ from previously characterized nisin-degrading proteases. Although candidate homologs of *ncr* and several proteases previously shown to cleave nisin in vitro were identified, all exhibited low sequence similarity to their respective reference proteins (Fig. S18 and Table S1). These findings suggest that nisin biotransformation in *B. stabilis* is mediated by previously uncharacterized extracellular proteases and highlight the potential for discovering additional enzymes involved in lanthipeptide turnover.

Finally, our results provide evidence that antimicrobial peptide production and biotransformation can occur within the same soil microbiome. The detection of both *nisA* and *nisZ* genes at the site from which the nisin-biotransforming isolates were recovered (Fig. 5) indicates the presence of multiple nisin-producing populations alongside bacteria capable of biotransforming the peptide. This coexistence suggests the potential for local nisin production and turnover within the soil environment. Rather than functioning solely as antimicrobial agents, lanthipeptides such as nisin may also contribute to nutrient cycling following their release into microbial communities. The extensive biotransformation observed in *P. fragi* (Fig. 2) and *B. stabilis* (Fig. 3), together with enhanced growth in nisin-supplemented media (Fig. 1B-C), supports the view that resistant environmental bacteria can derive nutritional benefit from antimicrobial peptides. These observations suggest that antimicrobial peptide resistance and antimicrobial peptide recycling may represent complementary ecological traits, linking survival during antimicrobial exposure with the subsequent recovery of peptide-derived nutrients. Collectively, our results expand the ecological framework of antimicrobial peptides, positioning them as dual-function molecules that contribute to both microbial antagonism and nutrient cycling within complex microbial communities.

## Materials and methods

### Culture media and reagents

M9 minimal medium was prepared by combining 200 mL of 5× M9 salts, 2 mL of 1 M MgSO₄, 100 µL of 1 M CaCl₂, and glucose, and adjusting the final volume to 1 L with Milli-Q water. Glucose concentrations were adjusted depending on the experiment: 0.01% for bacterial isolation, 0.01% or 0.05% for growth assays, and 0.5% for enzyme assays. King’s B (KB) medium was prepared with 5 g Bacto Proteose Peptone No. 3 (Gibco), 4.5 mL of 1 M K₂HPO₄, 9 mL of 1 M MgSO₄, and 10 mL of 50% glycerol, adjusted to a final volume of 1 L with Milli-Q water. KB medium was used for proteolytic assays. Lennox Broth (LB) (Carl Roth) was used for overnight bacterial cultures. For solid media, agar was added at a final concentration of 1.5%.

### Isolation of nisin-biotransforming bacteria from soil

Soil (1 g), collected from site P9 (55°47’19.7“N 12°33’29.9”E) in Jægersborg Deer Park (Dyrehaven), was suspended in 10 mL sterile phosphate-buffered saline (PBS), vortexed for 1 min, and allowed to settle for 2 min. An aliquot of the resulting suspension (100 µL) was inoculated into 50 mL sterile M9 minimal medium supplemented with 0.01% (w/v) glucose and 0.01% (w/v) nisin (nisin Z, with a purity greater than 95%, purchased from BOC Sciences, New York, USA). Cultures were incubated at 28 °C with shaking at 180 rpm for seven days. Following enrichment, 100 µL of the diluted culture was spread onto M9 agar plates containing either 0.01% (w/v) nisin or no nisin. Plates were incubated at 28 °C for 14 days. Colonies appearing on the nisin-containing plates were selected and streaked onto LB agar supplemented with 0.01% (w/v) nisin to obtain isolated colonies. Pure cultures were grown overnight and preserved in 20% glycerol stocks at −80 °C.

### Growth assays

Bacterial growth was monitored by measuring optical density at 600 nm (OD_600_). Overnight bacterial pellets grown in LB medium were harvested by centrifugation, washed three times with M9 minimal medium, and resuspended in M9 to an initial OD_600_ of 0.02. A sterile nisin stock solution (0.02%) was prepared in M9 medium.

Growth assays were conducted in sterile 96-well flat-bottom microplates (Greiner Bio-One). For each condition, 90 µL of the bacterial suspension was combined with 90 µL of either nisin-containing M9 medium or nisin-free M9 medium to generate the following treatments: (i) bacteria with nisin, (ii) nisin without bacteria, and (iii) bacteria without nisin. Plates were sealed with a sterile breathable membrane (Diversified Biotech, Breda, Netherlands) and incubated at 28 °C with continuous orbital shaking in a BioTek Cytation 5 plate reader. OD_600_ measurements were recorded every 10 min, and growth curves were analyzed using QurvE software. After incubation, culture mixtures were collected for downstream analyses. All experiments were performed in triplicate.

### LC-MS analysis

For chromatographic separation, 10 µL of each sample was loaded onto a Kinetex 2.6 µm polar C18 100 Å (100 x 2.1 mm). The analysis was performed at 0.4 mL/min and 40 °C on an Agilent 1290 Infinity II UPLC system under the gradient elution of 20 mM formic acid (FA) in water (A) and in methanol (B): 0–1 min: 2% B; 1–13 min: 2 to 70% B; 13–15.5 min: 70–100% B; 15.5–18.5 min: 100% B; 18.5–18.51 min: 100–2% B; 18.51–20 min: 2% B. All solvents were LC-MS grade and purchased from Sigma-Aldrich.

The analysis was carried out on a Bruker Tims TOF Flex (Bruker Daltonics, Bremen, Germany) instrument with electrospray ionization (ESI+) in a full scan mode with the following parameters: scan: 50 – 1400 *m/z*; end plate offset: 500 V; capillary: 3500 V; nebulizer: 3 bar; dry gas: 11 L/min; dry temperature: 250 °C; funnel 1 and 2 RF: 220 Vpp; multipole RF: 220 Vpp; collision energy and RF: 7 eV and 800 Vpp; transfer time: 75 µs; pre pulse storage: 8 µs. The analytical data were processed using Bruker Compass Data Analysis software version 6.1 (Bruker Daltonics GmbH & Co. KG).

### Identification of nisin biotransformation products

Culture mixtures obtained from the growth assays were analyzed by LC–MS to assess nisin abundance and identify potential biotransformation products. Nisin depletion in samples containing both bacteria and nisin relative to nisin-only controls was used as an indicator of nisin biotransformers.

For product extraction, cultures were transferred to 2 mL microcentrifuge tubes, and the microplate wells were rinsed with 200 µL of 0.2% formic acid (FA) in water, followed by 900 µL of 0.2% FA in methanol. The rinses were combined with the corresponding samples, centrifuged to remove cellular debris, and evaporated to dryness under nitrogen. The dried extracts were reconstituted in 360 µL of 0.2% FA in water before LC–MS analysis.

LC–MS data were processed to identify compounds present in cultures containing bacteria and nisin but absent in both bacteria-only and nisin-only controls. These compounds were considered candidate nisin biotransformation products. Structural assignments were proposed based on retention time, isotope distribution, exact mass and high-resolution mass accuracy, relative to the known structure of nisin. Selected products representing endpoint biotransformation products within the proposed pathways were further analyzed using MS/MS fragmentation to support structural assignments by elucidating product ion spectra.

### Cell-free supernatant preparation and extracellular enzyme assays

Bacterial pellets obtained from the overnight culture of strain 2 were washed three times and resuspended in sterile water. An aliquot (10 µL) of the suspension was inoculated into 60 mL of either M9 medium supplemented with 0.5% glucose or KB medium and incubated at 28 °C with shaking at 180 rpm for 70 h (three biological replicates). Cultures were then centrifuged and sterile-filtered to obtain cell-free supernatants. A portion of the cell-free supernatant was heat-treated at 100 °C for 15 min, cooled, and sterile-filtered to generate a heat-inactivated control. Separately, a sterile 0.03% nisin solution was prepared.

Extracellular enzyme assays were performed by mixing 2 mL of the untreated cell-free supernatant with 1 mL of the nisin solution. Control reactions consisted of either 2 mL of the heat-treated supernatant with 1 mL of nisin solution, or 2 mL of untreated supernatant with 1 mL of sterile water. Aliquots (500 µL) of each reaction were incubated at 28 °C with shaking at 180 rpm for 100 h for M9 and 70 h for KB. Reactions were quenched with 1.5 mL of 0.2% FA in methanol, after which the supernatants were evaporated under nitrogen and reconstituted in 1.5 mL of 0.2% FA in water prior to LC–MS analysis.

### Detection of nisin biosynthesis genes in soil microbiomes

Soil was collected from three sites in Dyrehaven Park, Denmark (P5: 55°47’19.7“N 12°33’29.9”E; P9: 55°47’28.2“N 12°34’30.4”E; P16: 55°47’40.5“N 12°34’19.6”E). At each site, six samples collected from a 10 cm depth were combined into a representative sample and stored at –80 °C. Tools were autoclaved and handling was performed in laminar flow cabinets to prevent contamination.

DNA was extracted using the DNeasy® PowerSoil® Pro Kit (Qiagen) and diluted to 50 ng/µL. Nisin primers (forward: 5’-GGACTATTCTTTAAACGCCTCGACGA-3’, reverse: 5’-TCTCAATGACTTGAGTATATTCAGTAAAACTCCG-3’) amplified a 791 bp fragment covering *nisA*/*nisZ* gene (174 bp) and part of *nisB* gene. PCR was performed with DreamTaq DNA Polymerase (Thermo) under the following conditions: 95 °C for 3 min; followed by 35 cycles of 95 °C for 30 s, 63.2 °C for 30 s, and 72 °C for 1 min; a final extension at 72 °C for 10 min. Products were separated on 1% agarose, bands (∼1000 bp) excised, purified (Macherey-Nagel protocol), and sequenced by Eurofins.

Sequences were analyzed with CLC Main Workbench v22 (Qiagen). Three replicates were analyzed per site.

### Genome sequencing and bioinformatic analyses

For DNA extraction, the nisin biotransformers were streaked onto LB agar plates. From these plates, individual colonies were harvested and cultured overnight in LB broth. A volume of 1 mL from the bacterial culture was used to extract DNA using the Monarch High Molecular Weight (HMW) DNA Extraction Kit (New England Biolabs, Germany).

DNA samples were processed using the Qubit dsDNA HS Kit to measure DNA concentration with the Qubit 3.0 Fluorometer (Invitrogen). The samples were then prepared for sequencing using the Rapid Barcoding Kit 24 V14 (SQK-RBK114.24) and sequenced on the GridION platform to at least 40X coverage. The resulting sequences were filtered for length (>1000bp) with nanoq (35), trimmed for adaptors with Filtlong (36) and then assembled with Flye (37). The full assemblies were then annotated with bakta (38) using the full database. Taxonomic identity was inferred by autoMLST2 (https://automlst2.cs.uni-tuebingen.de/) (39).

### Identification of candidate nisin-biotransforming enzymes

Putative nisin-biotransforming enzymes associated with the nisin-resistance protein (ncr) in *Lactococcus lactis* (A0A8A5P2H2) were identified by homology matching. Briefly, the ncr protein was blasted against the ClusteredNR database at NCBI, and all matches were downloaded and compiled into a custom blast database. Next, all predicted proteins from the isolates were blasted against this database. For confirmation, the process was repeated using an HMM approach with hmmsearch, and hits with E-values < 1.0e-09 were accepted. All protein hits and their matches were represented as a tree using the clustal-omega and simple-phylogeny pipeline at EMBL-EBI.

## Acknowledgements

We would like to acknowledge the DTU Metabolomics Core for use of LC-MS equipment. This study has received funding from the Danish National Research Foundation (DNRF137) for the Center for Microbial Secondary Metabolites (CeMiSt) and the Novo Nordisk Foundation (grant NNF24OC0093315).

